# Mastication while rested does not improve sustained attention in healthy participants conducting short-duration cognitive tasks

**DOI:** 10.1101/2024.02.12.579989

**Authors:** Devon Hansen, Greg Maislin, Jon E. L. Day

## Abstract

Sustained attention is important for optimal neurobehavioral performance, but many biological and environmental factors (e.g., circadian rhythm, distraction, etc.) may cause sustained attention deficits. It has been suggested that mastication (chewing) may ameliorate such deficits. As part of a continuing program to study the effects of mastication under varying conditions of fatigue and cognitive demand, this trial used a randomized, within-subjects, cross-over design to investigate the effect of mastication (gum chewing) on levels of sustained attention. To initially provide data that was ecologically valid for the average person, participants were not sleep deprived or otherwise challenged. Fifty-eight healthy adults (aged 18 – 45 years; 38 females) completed a 5 h in-laboratory daytime study during which time they completed two, 40 min test bouts. Participants completed the Psychomotor Vigilance Test (PVT), the Sustained Attention to Response Task (SART), the Karolinska Sleepiness Scale (KSS) and the Positive and Negative Affect Schedule (PANAS). During one of the two test bouts, participants were instructed to chew a piece of gum at a steady, comfortable rate. The statistical analyses were conducted blind. The primary outcome variable used for analyses was PVT lapses using the transformation square root of lapses plus square root of lapses plus 1 in addition to PVT mean reciprocal response time. Secondary outcome variables were PVT time-on-task slope and SART error score. Using rested participants and moderately fatiguing tasks, we were unable to detect any significant improvement in PVT or SART performance, or in KSS or PANAS ratings. A follow-up study under conditions of sleep deprivation and/or with longer task duration may provide further insight into the countermeasure potential of mastication.

## Introduction

Research into the effect of mastication on cognition started in the late 1930s when it was suggested that chewing sugar-coated chicle might help to manage stress levels and improve performance [1]. Despite the intensity of research into cognitive domains such as mood, memory and responses to stress over the last 16 years, it remains unclear precisely how mastication influences cognition [2-8] However, systematic reviews suggest that mastication can positively influence levels of alertness and sustained attention in vigilance tasks [9]. A more recent meta-analysis [10] detected a weak, but statistically significant improvement in levels of sustained attention when chewing. The meta-analysis also showed a tendency for feelings of alertness to decrease less during tasks requiring sustained levels of vigilance when chewing.

As part of a continuing program to study the effects of mastication under varying conditions of fatigue and cognitive demand, this trial used a randomized, within-subjects, cross-over design to investigate the effect of gum chewing on levels of sustained attention. To initially provide data that was ecologically valid for the average person, participants were not sleep deprived or otherwise challenged. Sustained attention was measured using tasks previously demonstrated to be sensitive in detecting any underlying fatigue.

The objective of the study was to characterize the impact of mastication (chewing gum) on sustained attention. The research hypothesis was that chewing improves measures of sustained attention. Specifically, it was predicted that mastication, as compared to non-mastication control, would improve performance by reducing performance instability and diminishing the time-on-task effect on the Psychomotor Vigilance Test (PVT) [11], and by reducing the error score on the Sustained Attention to Response Task (SART) [12].

## Materials and methods

### Participants and procedures

Thirty-eight healthy adults (38 women and 20 men) aged between 18 and 45 years (mean = 28.0 years; *SD* = 7.5 years) completed a 5 h highly controlled in-laboratory study. Participants were financially compensated for their study participation. They gave written informed consent and the study was approved by the Institutional Review Board (IRB) of Washington State University (WSU) and conformed with the Code of Ethics of the World Medical Association (Declaration of Helsinki).

Prospective participants were initially pre-screened over the telephone. If eligible and available, they were scheduled for an in-office screening session at the Sleep and Performance Research Center (SPRC) at WSU Spokane. They were in the age range of 18 - 45 years. They were physically and psychologically healthy. Participants had no sleep or circadian disorders, as confirmed with validated questionnaires and clinical interview by a designated medical professional for the study. Participants were free of drugs and alcohol, as assessed by physical examination, urine drug screen, and alcohol breath test. They had no clinically relevant history of psychiatric illness and no clinically relevant history of brain injury. They were not current tobacco users and reported no current medical or drug treatment (excluding oral contraceptives). They were not vision impaired or their vision was corrected to normal. Their habitual sleep duration was between 6 and 10 h per day.

Participants were asked to maintain their habitual sleep schedule (within ± 30 min) in the three days prior to the experiment with strict regularity as verified by a sleep diary and messages left on a time-stamped voicemail box. Further, in the three days prior to their experiment, sleep was monitored by means of actigraphy to measure their sleep/wake activity patterns.

The study utilized a within-subjects cross-over design in which participants completed both conditions (mastication vs. control). Order of chewing-gum administration was randomized and counterbalanced across the two testing blocks.

The study took place inside the Human Sleep and Cognition Laboratory of the SPRC at WSU Health Sciences in Spokane, WA, USA. Participants were studied in groups of up to four and were assigned their own room for performance testing. The laboratory environment was carefully controlled, with a fixed ambient temperature (22 ± 1 °C), and fixed light levels (< 100 lux). Participants were not allowed to use phones, computers, or otherwise communicate with the outside world during their time in the laboratory. Participants were asked to refrain from consuming caffeine, alcohol, tobacco, and chewing gum (beyond what was provided during testing).

Participants reported to the laboratory at 09:00. Following study admission procedures and training on performance tasks, participants had a 1 h break before beginning the first of two test bouts at 11:00. Each test bout was approximately 40 min long. In between test bouts, subjects had a 1 h break inside the laboratory. During one of the two test bouts, subjects were instructed to chew a piece of gum at a steady, comfortable rate. Half of the sample was assigned to the mastication condition during the first test bout, the other half during the second test bout (Figure 1).

**Figure 1:**
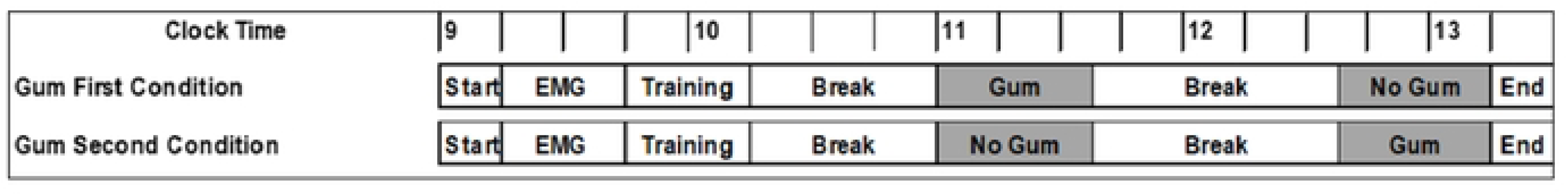
*schematic of the study design by mastication condition. Clock time is indicated at the top of the figure; test bouts are shaded in grey*.

Study admission included review of the participant’s sleep diary and wrist actigraphy to verify bedtimes and wake time were consistent (± 30 min from their self-reported habitual times on average). Habitual reported bedtime was 22:50 (*SD* = 1.0 h); habitual reported wake time was 07:11 (*SD* = 1.2 h). Participants also completed a urine drug screen and alcohol breathalyzer test to ensure they were drug and alcohol free. Following admission procedures, participants were fitted with equipment for electromyography (EMG; Nihon Kohden, Irvine, CA) to verify mastication activity during subsequent testing. The EMG montage included submental (T3, T4) and masseter.

### Mastication intervention

Spearmint flavoured chewing gum (Extra Brand, Wrigley, Inc) was administered during one (of two) testing session. Prior to each study run, each stick of gum was unwrapped to shield branding information and placed individually in a clear plastic bag each labelled with the date, participant ID number, and gum administration order. Immediately prior to the start of the test bout with gum administration, participants were provided with their labelled gum and instructed to chew at a comfortable but steady pace throughout the testing session. Chewing activity was verified via EMG recordings. If chewing stopped during testing, participants were instructed to continue chewing via an intercom system.

### Neurobehavioral testing

The cognitive test battery included the SART [12] and the PVT [11]. The primary outcome variable used for analyses was PVT lapses (defined at reaction times > 500 ms) using the transformation square root of lapses plus square root of lapses plus 1 in addition to PVT mean reciprocal response time as this has been shown to be a more sensitive metric for detecting treatment effects. Secondary outcome variables were PVT time-on-task slope and SART error score.

In their assigned room, participants were trained on the performance tasks by a trained staff member. The test battery included, in order, completion of a subjective rating of mood (Positive and Negative Affect Scale; PANAS) and fatigue (Karolinska Sleepiness Scale; KSS), a 20-min version of the Sustained Attention to Response Task (SART), a 10-min version of the psychomotor vigilance test (PVT) and concluded with completion of the PANAS and KSS again.

### Psychomotor Vigilance Test (PVT)

The PVT requires participants to respond to a visual stimulus (incrementing millisecond counter) by pressing a response button [11]. The fore-period prior to stimulus presentation was randomized between 2 and 10 s in 1 s increments. Participants were instructed to respond as quickly as possible as soon as the stimulus appeared but not to respond before the stimulus appeared. Following each response, the response time (RT) was displayed (in ms) for 1 s. Performance was quantified as the number of lapses (RTs > 500 ms) and false starts (RTs < 150 ms).

### Sustained Attention to Response Task (SART)

The SART is a go/no go task [12]. Participants were required to either respond to or withhold a response to a visual stimulus by pressing the space bar on a computer keyboard. The fore-period prior to stimulus presentation was fixed at 1150 ms and each stimulus was presented for 250 ms. Participants were instructed to respond by pressing the space bar to every stimulus presented as quickly as possible but were to withhold their response when a specific numeric stimulus appeared. Stimuli were presented randomly in five different font sizes. Performance was quantified as the numbers of errors (incorrect space bar presses). Two different versions with unique stimuli of the SART were administered in counterbalanced order.

Immediately following a training session, participants had a 1 h break before commencing testing at 11:00. Following the first 40 min test bout, participants had a 1 h break. During the break, participants were allowed to engage in low-stimulation activities such as reading, watching a movie, playing a board game, or talking with the other participants or staff members, but they could not leave the laboratory. Following the 1 h break, participants completed their second 40 min test bout. Immediately following the second test bout, the EMG electrodes were removed, and participants were asked to complete an end of study questionnaire before leaving the laboratory.

### Subjective ratings (PANAS and KSS)

Subjective ratings of sleepiness were taken using a computerized version of the Karolinska Sleepiness Scale (KSS) [13]. This is a Likert-type rating scale on which subjects rated their sleepiness, ranging from 1 (very alert) to 9 (very sleepy). Participants also completed the Positive and Negative Affect Schedule (PANAS) [14], a 20-item questionnaire asking participants to self-rate the current applicability of each of 20 affective words that describe feelings and emotions using a scale from 1 (“very slightly or not at all applicable”) to 5 (“extremely applicable”).

### Power analysis

The SART was used to determine sample size. This decision was based on factors: 1) that there was prior literature in this area; 2) that it was predicted that the SART would exhibit a smaller effect size than the PVT. The power calculation was based on data taken from previous work [15] in which 20 participants reported correct mean performance scores on the SART of 10.75 and 7.77 (mastication vs. control) with standard error of the mean that did not exceed 2 (see Figure 1 in [15]). This implied an effect size of at least 0.33. With a one-sided type 1 error threshold of α = 0.05 and power of 80% to detect performance changes between mastication and control, the minimum sample size for this effect size was *n* = 58 subjects (nQuery Advisor 7.0).

### Statistical methods for the analysis of primary and secondary outcome measures

The initial data analyses for PVT, SART, KSS and PANAS were conducted blinded regarding experimental condition (mastication or control) which were designated as conditions “A” and “B”. There were two treatment sequences, A then B and B then A.

The primary outcome variable used for analyses was PVT lapses using the transformation square root of lapses plus square root of lapses plus 1 in addition to PVT mean reciprocal response time as this has been shown to be a more sensitive metric for detecting treatment effects. The transformation used was the Freeman-Tukey transformation [16] for Poisson random variable which tends to normalize the distribution of PVT lapses as well as stabilizes the variance. Secondary outcome variables were PVT time-on-task slope and SART error score.

Descriptive summary statistics were provided which included the sample size, mean, standard deviation, standard error, minimum, and maximum, for each of the four cells defining the cross-over design, the treatment difference (mastication vs. control) within each treatment sequence, the overall treatment difference, and the overall period difference.

The standard deviation reported for a “difference” was determined as the pooled standard deviation across both treatment sequences (A/B and B/A), assuming equal variances. The standard error is the standard deviation of the mean estimate.

To determine any difference between treatment effect (mastication vs. control) and period (1 and 2), *t*-tests were performed using both pooled and Satterthwaite versions of the tests. The sample size analysis was based on a one-sided type 1 error rate of 0.05. However, there was no way to produce one-sided *p*-values when conditions were blinded. Therefore, two-sided *p*-values for mastication vs. control differences were summarized. When the data was unblinded, if the mastication minus control difference for a particular endpoint was in the hypothesized direction, the one-sided *p*-value was obtained by dividing the two-sided *p*-values by two. If the mastication minus control difference was in the opposite direction, the one-sided *p*-value is > 0.50. 95% two-sided confidence intervals were provided for statistical estimation purposes. Both pooled (assuming equal variances for both treatment sequences) and Satterthwaite (assuming unequal variances) intervals were provided. The folded *F*-test of equal variances in each treatment sequence were evaluated.

Period effects were evaluated. Period effects may arise when subjects do better in a subsequent period because their state has changed, for example, their mental or health status has changed, independent of treatment.

The presence of baseline differences in the subjective ratings (sessions 3 and 5) was first assessed using one-way analysis of variance. If significant differences at baseline were detected, the baseline was subtracted from the subjective ratings in sessions 4 and 6 respectively to obtain a change score. The data were then analyzed using a mixed-model analysis of variance (ANOVA). Fixed effects for chewing intervention (within-subject factor: ‘chewing’ or ‘not chewing’), sequence (between-subject factor: either A (chewing) then B (not chewing) or (not chewing) then (chewing), and intervention-by-sequence interaction were included. Planned contrasts were included for pairwise comparisons between interventions overall and in interaction with sequence. Bonferroni correction was used to adjust for the effects of multiple comparisons. There was one missing rating in the PANAS ratings for one subject (session 3).

## Results

### Analysis of primary and secondary outcome measures

#### PVT lapses

The treatment effect (mastication vs. control) for sequences 1 and 2 were 0.055 (95% CI [-0.304, 0.414]) and 0.129 (95% CI [-0.373, 0.633]). Assuming equal standard deviations between sequences, the pooled treatment effect over sequences (chewing vs. non-chewing) was 0.093 (95% CI [-0.210, 0.394]). Although there was some evidence of a difference between sequences in the standard deviations of treatment differences, *F*_(28,28)_ = 1.97, *p* = 0.079, the confidence interval and *t*-test results for the treatment effect using the Satterthwaite approach were nearly identical (results not shown). The pooled *t*-test value was 0.61 with 56 degrees-of-freedom. The 2-sided *p*-value for superiority is 0.54. Since the results were in the opposite direction from what was hypothesized, the 1-sided *p*-value > 0.50. The within subject standardized effect size was 0.093/0.574 = 0.162 which is conventionally considered small [17]. Therefore, the quantitative analysis of transformed lapses does not allow for rejecting the null hypothesis given this sample size.

**Table 1:** results summary according to experimental cell for transformed PVT lapses.

|  | Period 1 |  |  |  | Period 2 |  |  |  |
| --- | --- | --- | --- | --- | --- | --- | --- | --- |
| | Intervention | Mean | SD | $n$ | Intervention | Mean | SD | $n$ |
| Sequence 1 | Mastication | 1.981 | 1.072 | 29 | Control | 1.926 | 1.147 | 29 |
| Sequence 2 | Control | 1.858 | 1.290 | 29 | Mastication | 1.988 | 1.475 | 29 |

#### SART

The treatment effect (mastication vs. control) for sequences 1 and 2 were -0.035 (95% CI [-1.334, 1.265]) and 0.690 (95% CI [-1.066, 2.445]). Assuming equal standard deviations between sequences, the pooled treatment effect over sequences (mastication vs. control) was 0.328 (95% CI [-0.740.1.396]). The standard deviation of differences did not significantly differ between sequences, *F*_(28,28)_ = 1.82, *p* = 0.118. The pooled *t*-test value was also 0.61 with 56 degrees-of-freedom (2-sided *p*-value for superiority = 0.54). Since the results were in the opposite direction from what was hypothesized, the 1-sided *p*-value > 0.50. The within subject standardized effect size was 0.328/2.030 = 0.162 which is conventionally considered small. Therefore, the quantitative analysis of SART errors does not allow for rejecting the null hypothesis given this sample size.

**Table 2:**
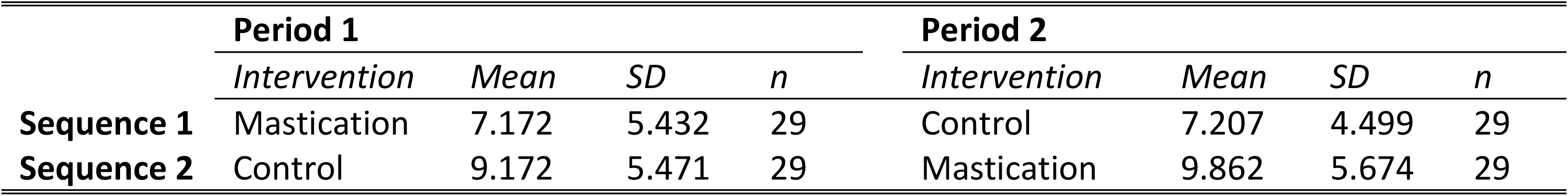
results summary according to experimental cell for SART errors.

**Table 3:**
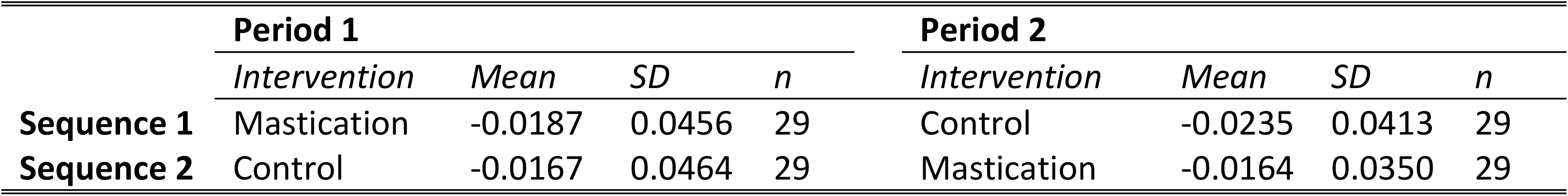
results summary according to experimental cell for PVT time on task slope.

|  | Period 1 |  |  |  | Period 2 |  |  |  |
| --- | --- | --- | --- | --- | --- | --- | --- | --- |
| | Intervention | Mean | SD | $n$ | Intervention | Mean | SD | $n$ |
| Sequence 1 | Mastication | -0.0187 | 0.0456 | 29 | Control | -0.0235 | 0.0413 | 29 |
| Sequence 2 | Control | -0.0167 | 0.0464 | 29 | Mastication | -0.0164 | 0.0350 | 29 |

#### PVT time-on-task slope

The treatment effect (mastication vs. control) for sequences 1 and 2 were 0.0048 (95% CI [-0.0150, 0.0247]) and 0.000268 (95% CI [-0.0198, 0.0204]). Assuming equal standard deviations between sequences, the pooled treatment effect over sequences (mastication vs. control) was 0.00254 (95% CI [-0.0113, 0.0164]). The standard deviation of differences did not significantly differ between sequences, *F*_(28,28)_ = 1.02, *p* = 0.951. The pooled *t*-test value was 0.37 with 56 degrees-of-freedom (2-sided *p*-value for superiority = 0.71). Since the results were in the expected direction, the 1-sided *p*-value is 0.36. The within subject standardized effect size was 0.00254/0.0263 = 0.097 which is conventionally considered small. Therefore, the quantitative analysis of PVT Time on Task errors does not allow for rejecting the null hypothesis given this sample size.

The objective of the experimental study was to investigate the effect of mastication on levels of sustained attention measured in PVT and SART computerized tasks. The blind analyses found no evidence in support of the claims. Using this test paradigm, no significant improvements in PVT or SART performance were detected.

There were no significant mastication vs. control condition effects for either the primary endpoints (PVT lapses) or secondary endpoints (SART total errors and PVT reciprocal time on task slope).

However, the results for PVT lapses are challenged due to the generally lower prevalence of response time lapses among normal healthy volunteers. Before unblinding, a supplemental endpoint was added due to its known greater sensitivity to detect differences in a variety of experimental settings, PVT mean reciprocal response time.

Following unblinding, it was determined that the primary null hypothesis concerning PVT lapses could not be rejected (one-sided *p* > 0.50) whether based on transformed lapses or untransformed lapses. The mean difference was not significantly different from zero. The mean difference was slightly positive and not negative as predicted by the research hypothesis. It did not appear that the mean difference was significantly associated with age or average performance level.

The secondary null hypothesis concerning SART errors could not be rejected (one-sided *p* > 0.50). The mean difference was not significantly different from zero. The mean difference was slightly positive and not negative as predicted by the research hypothesis. It did not appear that the mean difference was significantly associated with age or average performance level.

The secondary null hypothesis concerning PVT reciprocal RT time on task slope could not be rejected (one-sided *p* = 0.36). The mean difference was not significantly different from zero. The mean difference was slightly positive. Therefore, the direction of the mean difference was the direction predicted by the research hypothesis. This was the only outcome variable in which the direction of the mean difference was in the direction predicted by the research hypothesis. It did not appear that the mean difference was significantly associated with age or average performance level.

#### Subjective ratings

No significant effects of sequence were detected in baseline sessions 3 and 5. Therefore, baseline subtraction was not performed and the KSS and PANAS ratings in sessions 4 and 6 were analysed directly. A significant interaction between intervention and sequence was detected for both KSS (*F*_(1,56)_ = 5.304, *p* < 0.05) and PosPANAS (*F*_(1,56)_ = 15.306, *p* < 0.001) but not NegPANAS (NS). However, no significant differences were detectable in the *post hoc* comparisons of means for either KSS or PosPANAS following Bonferroni correction (results not shown).

## Discussion

The objective of this study was to characterize the impact of mastication (chewing gum) on sustained attention. The research hypothesis was that mastication improves measures of sustained attention. Specifically, it was predicted that mastication, as compared to a non-mastication control, would improve performance by reducing performance instability and attenuating the vigilance decrement (intervention x time-on-task effect) on the PVT, and by reducing the error score on the SART.

Contrary to our predictions, no significant improvements in SART or PVT performance were detected. This suggests that either: 1) mastication does not significantly improve levels of sustained attention; 2) the testing duration was too short and participants did not become sufficiently fatigued to enable mastication to sustain levels of attention, and 3) the presence of baseline differences increased the signal to noise ratio (masking [potentially small] treatment effects).

Sustained attention tasks are intended to induce a vigilance decrement. Effective interventions would be expected to maintain levels of performance over time. Some previous studies have reported significant intervention x time-on-task effects suggesting that performance declines less rapidly over time when chewing compared to when not chewing [18-20]. A longer test duration would be expected to impose a more pronounced decrease in sustained attention and therefore a greater chance of observing a significant treatment x time interaction.

This study examined the effects of mastication in an ecologically valid (unchallenged, normal, everyday) situation. While it is possible that the effects of chewing on levels of sustained attention may only be a weak effect, further blind clinical trials need to determine if an effect of mastication on sustained attention can be demonstrated. Improvements might become detectible by testing under conditions of sleep deprivation and/or with longer task duration or in a time of the day where the effect of fatigue or low-energy is more pronounced. Furthermore, additional insights into the potential of mastication to improve levels of sustained attention might be accessed by varying chewing intensity through gum texture and flavour, rate of chewing and temporal relationship of chewing relative to the sustained attention task.

## Acknowledgements

Sophie Miquel for her contribution to the design of the trial.

